# The RNA-binding protein HuR promotes nonalcoholic steatohepatitis (NASH) progression by enhancing death signaling pathway

**DOI:** 10.1101/2020.10.21.348185

**Authors:** Xinzhi Li, Bingchuan Yuan, Zhicheng Yao, Xu Sun, Weixiang Guo, Zheng Chen

## Abstract

Hepatocyte death triggers liver inflammation, liver injury, and fibrosis, which contributes to non-alcoholic steatohepatitis (NASH) pathogenesis. However, whether RNA processing regulates death signaling pathway during NASH progression is not investigated. In this study, we show that HuR, a widely expressed RNA-binding protein, promotes NASH progression by increasing DR5/caspase8/caspase3-mediated hepatocyte death. Cytosolic HuR levels are abnormally elevated in human patients with NASH. Hepatocyte-specific deletion of *HuR* protects against MCD-induced NASH by decreasing liver steatosis, inflammation and cell death, whereas hepatic overexpression of HuR induces liver injury by increasing DR5-induced hepatocyte death. Furthermore, in primary hepatocytes, HuR deficiency ameliorates PA&TNFα-induced hepatocyte death due to decreased DR5/caspase8/caspase 3 signaling pathway while overexpression of HuR induces hepatocyte death by increasing DR5/caspase8/caspase 3 signaling pathway. Mechanistically, HuR directly binds to 3′-UTR of DR5 transcript and promotes its mRNA stability, contributing to the hepatocyte death during NASH progression. Our data reveal a novel mechanism by which HuR promotes mRNA stability of DR5, which contributes to NASH progression.

## Introduction

Non-alcoholic steatohepatitis (NASH) is characterized by hepatic steatosis, liver injury, chronic inflammation and liver fibrosis, which has been recognized as a key step for the development of cirrhosis and hepatocellular carcinoma (HCC) ^1, 2^. A ‘two hit’ theory has been proposed to explain NASH pathogenesis ^3^. The first hit is hepatic steatosis due to abnormal lipid accumulation in the liver ^4, 5^. The second hits include oxidative stress, inflammation, cell death and liver injury ^3, 6, 7^. Hepatocyte death is at the center of the second hits because it triggers liver inflammation, liver injury, and fibrosis. Clinical and genetic mouse studies demonstrate that hepatocyte apoptosis plays a key role in NASH progression^8, 9, 10, 11, 12^. It has been shown that the ligands for death receptors (e.g., TNFα and TRAIL), death receptors (e.g., FAS and DR5), and the activation of caspases 2, 3, and 8 are abnormally upregulated in the liver of human patients with NASH ^11, 12, 13, 14, 15^. TRAIL knockout mice are protected from diet-induced NASH ^16^. The free fatty acid such as palmitate can induce aggregation of DR5 on the cell membrane and then activate caspase 8/caspase 3 signaling pathway, leading to hepatocyte death^9^. Knockout of caspase 2 or hepatic deletion of caspase 8 attenuates methionine/choline deficient diet (MCD)-induced NASH ^12, 15^. The treatment of caspase inhibitors have been shown to protect against NASH pathogenesis^17, 18^. However, whether RNA processing regulates death signaling pathway during NASH progression is not investigated.

ELAV (embryonic lethal/abnormal visual system)/Hu family RNA-binding proteins, such as ELAVL1/HuR/HuA, ELAVL2/HuB, ELAVL3/HuC and ELAVL4/HuD, contain three conserved RNA recognition motifs and specifically bind adenylate-uridylate-rich element (AREs) in the 3′-untranslated region (UTR) of targeted mRNAs, regulating RNA stability and translational efficiency ^19, 20, 21, 22^. HuB, HuC and HuD are exclusively expressed in neurons, whereas HuR is ubiquitously expressed, including liver ^21^. Several studies have shown that HuR play important roles in the liver de-differentiation, liver development, human hepatocellular carcinoma (HCC) progression, and liver fibrosis ^23, 24, 25^. However, whether HuR regulates NASH progression by modulating RNA processing of death signaling molecules is still largely unknown.

In this study, we show that abnormally elevated expression of HuR promotes the pathogenesis of NASH. Hepatocyte-specific deletion of *HuR* protects against NASH progression by decreasing liver steatosis, inflammation and cell death, whereas hepatic overexpression of HuR induces liver injury by increasing DR5/caspase8/caspase3-induced cell apoptosis. Furthermore, HuR directly binds to 3′-UTR of DR5 transcript and promotes its mRNA stability, contributing to the hepatocyte death during NASH progression. This study also suggests that HuR might be a novel drug target for the treatment of NASH.

## Materials and Methods

### Animal Experiments

Animal experiments were carried out in strict accordance with the Guide for the Care and Use of Laboratory Animals published by the US National Institutes of Health (NIH publication No. 85-23, revised 1996). Animal experiment protocols were approved by the Institutional Animal Care and Use Committee or Animal Experimental Ethics Committee of Harbin Institute of Technology (HIT/IACUC). The approval number is IACUC-2018004. Mice were housed on a 12-h light/12-h dark cycle with a free access to water. For MCD-induced NASH, mice were fed with an MCD (Medicience) for three weeks. Hepatocyte-specific *HuR* knockout mice were generated by crossing *HuR*^flox/flox^ mice ^26, 27^ with Alb-Cre mice. Blood samples were collected from orbital sinus. The serum alanine aminotransferase (ALT) activity and TAG levels was measured with an ALT or TAG reagent set, respectively ^28^.

### Human samples

Human liver samples were collected from the Third Affiliated Hospital of Sun Yat-sen University. The present study was approved by the Research Ethics Committee of the Third Affiliated Hospital of Sun Yat-sen University, and individual permission was obtained using standard informed consent procedures. The investigation conforms to the principles that are outlined in the Declaration of Helsinki regarding the use of human tissues. Samples with a NASH activity score (NAS) ≥ 5 or a NAS of 3–4 but with fibrosis were included in the NASH group and samples with a NAS of 0 were classified as non-steatotic control.

### Primary hepatocyte culture and Aadenoviral infection

Primary hepatocytes were isolated from C57BL/6 WT, *HuR*^flox/flox^ and *HuR-*HKO mice by liver perfusion with type II collagenase (Worthington Biochem, Lakewood, NJ) and cultured at 37°C and 5% CO2 in DMEM medium supplemented with 5% FBS. Primary hepatocytes from from C57BL/6 WT mice were infected with βGal or Flag-HuR adenoviruses overnight.

### Nuclear extract preparation

Liver tissues were homogenized in a lysis buffer (20 mM HEPES, 1 mM EDTA, 250 mM sucrose, 1 mM PMSF, 1 mM Na_3_VO_3_ and 0.5 mM DTT, pH 7.4) and centrifuged sequentially at 1,100g and 4,000g at 4 °C. Nuclear protein was extracted from the pellets using a high-salt solution (20 mM HEPES, 420 mM NaCl, 0.2 mM EDTA, 0.5 mM DTT, 1 mM PMSF and 1 mM Na_3_VO_3_, pH 7.9). The preparation of nuclear and cytosolic protein from primary hepatocytes was using a commercial Nuclear and Cytoplasmic Protein Extraction Kit (P0027, Beyotime).

### Immunoblotting and RNA-immunoprecipitation

For immunoblotting, total cell lysates, cytosolic and nuclear lysates were immublotted with the indicated antibodies, and visualized using the ECL as shown previously ^29^.

Primary hepatocytes were used for RNA-immunoprecipitation. Primary hepatocytes from C57BL/6 WT mice were infected with βGal or Flag-HuR adenoviruses overnight. Primary hepatocytes were also isolated from *HuR*^flox/flox^ and *HuR-*HKO mice and cultured overnight. Cell lysates from the above hepatocytes were immunoprecipitated with Flag beads and HuR antibody at 4 °C for 2 hours. The subsequential RNA immunoprecipitation (RIP) was performed using the Magna RIP kit (Merck Millipore) according to the manufacturer’s instruction. RNA samples retrieved from anti-HuR or Flag-beads were used for RT-qPCR. Antibody information was shown in Supplementary Table 1. Primers for qPCR was listed in Supplementary Table 2.

### Real Time Quantitative PCR (RT-qPCR)

RT-qPCR was performed as shown previously ^29, 30^. Briefly, total RNAs were extracted using TriPure Isolation Reagent (Roche, Mannheim, Germany), and the first-strand cDNAs were synthesized using random primers and M-MLV reverse transcriptase (Promega, Madison, WI). qPCR was done by using Roche LightCycler 480 real-time PCR system (Roche, Mannheim, Germany). The expression of individual genes was normalized to the expression of 36B4 (a house-keeping gene). Primers for real time qRT-PCR were listed in Supplementary Table 2.

### RNA sequencing

RNA-seq were performed as described previously ^29, 31^. The hepatic mRNA profiles of *HuR*^flox/flox^ and hepatocyte-specific HuR knockout (HKO) mice (n=3 for each group) fed with an MCD for 3 weeks were generated by deep sequencing using an Illumina Novaseq platform. The mRNA profiles of β-Gal-overexpressing, and HuR-overexpressing primary hepatocytes were also generated by deep sequencing using an Illumina Novaseq 6000 platform. Paired-end clean reads were aligned to the mouse reference genome(GRCm38.p6) with Hisat2 v2.0.5, and the aligned reads were used to quantify mRNA expression by using featureCounts v1.5.0-p3. RNA-seq data that support the findings of this study have been deposited in GEO under accession code GSE152670 and GSE154015.

### RNA decay assay

Primary hepatocytes from C57BL/6 WT mice were infected with βGal or Flag-HuR adenoviruses overnight. Primary hepatocytes were also isolated from *HuR*^flox/flox^ and *HuR-*HKO mice. Hepatocytes were treated with actinomycin D (10 mg/ml) for 0, 3, 6, and 12 h. Cells was harvested at indicated time, and RNA was isolated for RT-qPCR. The degradation rate of RNA (*k*) was estimated by plotting N_t_/N_0_ against time and fitting to the following equation^32^:

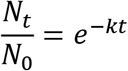

The half life of mRNA was calculated following this equation:

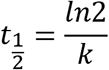

### Statistical Analysis

Data were presented as means ± S.E. Differences between groups were analyzed by Student's *t* tests. p< 0.05 was considered statistically significant. *, p< 0.05. **, p< 0.01.

## Results

### Cytosolic HuR is abnormally elevated in NASH livers

Cytosolic translocation of HuR has been shown to regulate RNA stability ^33, 34^. To test whether the cytosolic translocation of HuR happens in NASH, we measured cytosolic, nuclear and total HuR protein levels in NASH by immunoblotting. As shown in Fig. 1a, b, cytosolic and total HuR protein levels were significantly increased in the livers of human patients with NASH, whereas the nuclear HuR protein levels were not altered in NASH (Fig. 1a,b). Consistently, in mice with MCD-induced NASH, cytosolic and total HuR protein levels were also dramatically increased, whereas the nuclear HuR protein levels were not altered (Fig. 1c,d). To further determine how HuR expression was increased in NASH, primary hepatocytes were isolated and treated with TNFα or palmitate acid (PA), as it is known that TNFα and PA could mimic inflammation and lipotoxicity in NASH, respectively ^35, 36^. As shown in Figure 1e-h, both TNFα and PA could significantly increase the HuR protein levels in both cytosol and total lysates, whereas the nuclear HuR levels were not changed under these two conditions. These data indicate that inflammation and lipotoxicity could increase both HuR expression and cytosolic localization of HuR, which may further promote NASH.

**Figure 1.**
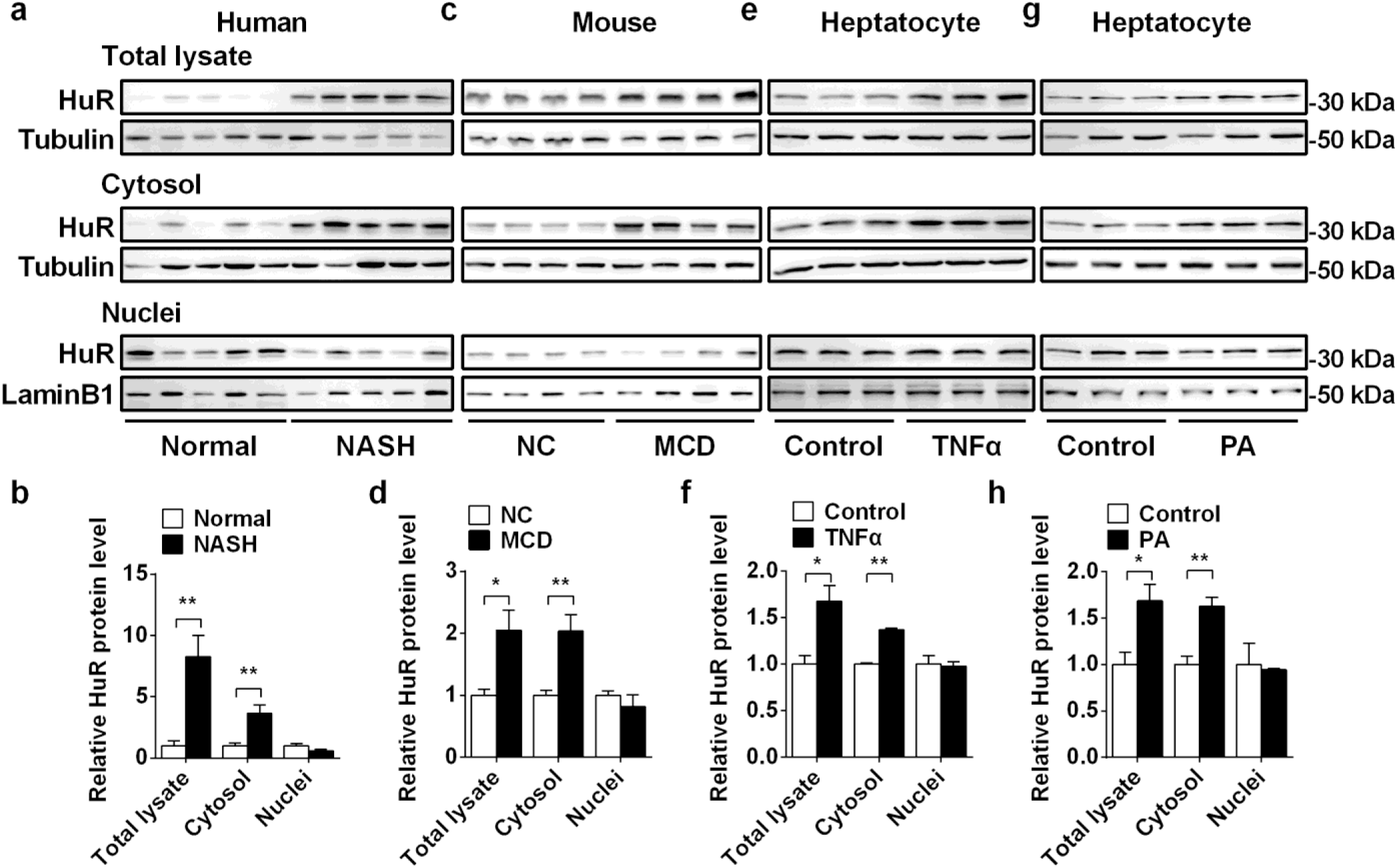
Cytosolic HuR is abnormally elevated in the NASH livers. (a) HuR protein levels in cytosol, nuclei, and total cell lysate from human NASH and normal liver tissues were measured by immunoblotting. (b) Quantification of the data in (a). (c) HuR protein levels in cytosol, nuclei, and total cell lysate from the livers of MCD (3 weeks)-fed mice were measured by immunoblotting. (d) Quantification of the data in (c). (e)Primary hepatocytes were isolated from C57BL6 mice and treated with TNFα (40 ng/ml) for 2 hours. HuR protein levels in cytosol, nuclei, and total cell lysate were measured by immunoblotting. (f) Quantification of the data in (e). (h) Primary hepatocytes were isolated and treated with palmitate acid (0.5 mM) for 18 hours. HuR protein levels in cytosol, nuclei, and total cell lysate were measured by immunoblotting. (h) Quantification of the data in (g). n=3-5 for each group. Each experiment was repeated at least twice with similar results. *, p< 0.05. **, p< 0.01. Data represent the mean ± SEM.

### Hepatic deletion of *HuR* protects against MCD-induced NASH

To determine whether HuR regulates the NASH progression, we generated hepatocyte-specific *HuR* knockout (HKO) mice by cross *HuR*^flox/flox^ mice with *Alb*-Cre transgenic mice. The genotype of *HuR*-HKO mice is *HuR*^flox/flox^ Alb-Cre^+/−^. As expected, HuR protein levels were dramatically decreased by 96.2% in the livers of *HuR*-HKO mice (Fig. 2a). We did not observe any difference between *HuR*^flox/flox^ and *Alb*-Cre mice (data not shown). *HuR*^flox/flox^ mice also displayed similar MCD-induced NASH with *Alb*-Cre mice, as revealed by similar serum ALT activity (Supplementary Fig. 1a), and similar liver TAG levels (Supplementary Fig. 1b). Therefore, we used *HuR*^flox/flox^ mice as the control for *HuR*-HKO mice in the following experiments. Under a normal chow diet feeding condition, the liver weights, liver TAG levels, and serum ALT activity were comparable between *HuR*-HKO and *HuR*^flox/flox^ mice (Fig. 2b-d). To further test whether HuR is involved in NASH progression, *HuR*-HKO and *HuR*^flox/flox^ mice were fed with an MCD for 3 weeks. *HuR*-HKO mice protected against MCD-induced NASH, as revealed by lower serum ALT activities (Fig. 2e), lower liver weights and sizes (Fig. 2f,g), less hepatic lipid droplets (Fig. 2h,i) and lower liver TAG levels (Fig. 2j) compared with *HuR*^flox/flox^ mice. We then measured hepatocyte death by performing TUNEL assays in MCD feeding mice. The TUNEL-positive cells were significantly decreased in MCD feeding *HuR*-HKO mice (Fig. 2k). Death receptors (such as DR5 and FAS)/caspase8 signaling pathway has been shown to regulate hepatocyte death ^8, 37^. Downregulation of this signaling pathway may be associated with decreased hepatocyte death in *HuR*-HKO mice. To test this hypothesis, the expression of death receptors and the cleavage of caspase8 and caspase 3 were measured by immunoblotting. As shown in Fig. 2l, the protein levels of DR5 and the cleavage of caspase8 were much lower in the liver of *HuR*-HKO mice whereas the FAS protein levels were comparable between *HuR*-HKO and *HuR*^flox/flox^ mice, which indicates that downregulation of DR5/caspase8 signaling pathway contributes to decreased hepatocyte death in *HuR*-HKO mice fed with MCD. Furthermore, liver fibrosis in MCD-fed *HuR*-HKO mice was ameliorated as revealed by less Sirius Red-positive areas in the liver sections than that in *HuR*^flox/flox^ mice (Fig. 2m). These data suggest that hepatic deletion of *HuR* protects against MCD-induced NASH.

**Figure 2.**
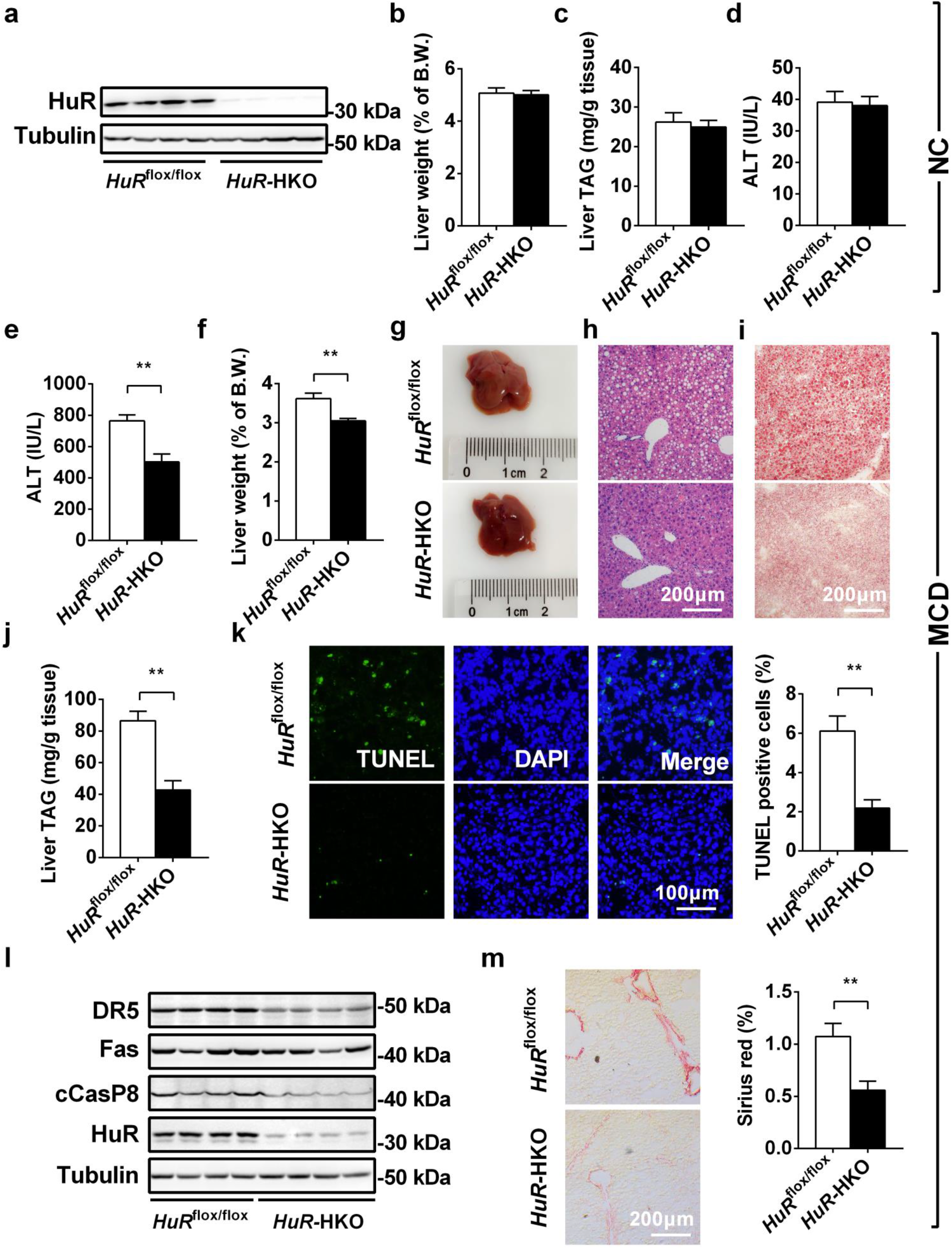
Hepatic Deletion of *HuR* ameliorates MCD-induced NASH. (a) HuR protein levels in livers of *HuR*^flox/flox^ and *HuR-*HKO mice at 8 weeks age (n=4). (b-d) Liver weight, liver TAG levels, and serum ALT activity in *HuR*^flox/flox^ and *HuR-*HKO mice fed with an NC at 8 weeks age (n=6-10). (e) Serum ALT activity in *HuR*^flox/flox^ and *HuR-*HKO mice fed with an MCD for 3 weeks (n=8-11). (f) Liver weight of *HuR*^flox/flox^ and *HuR-*HKO mice fed with an MCD for 3 weeks (n=7-9). (g) Representative liver pictures from *HuR*^flox/flox^ and *HuR-*HKO mice fed with an MCD for 3 weeks. (h, i) Representative H&E and Oil Red O staining of liver sections from *HuR*^flox/flox^ and *HuR-*HKO mice fed with an MCD for 3 weeks. (j) Liver TAG levels in *HuR*^flox/flox^ and *HuR-*HKO mice fed with an MCD for 3 weeks (n=7-9). (k) The TUNEL-positive cells in *HuR*^flox/flox^ and *HuR-*HKO mice fed with an MCD for 3 weeks (n=5). (l) The death receptors (FAS and DR5) /caspase8/caspase3 signaling pathway was measured by immunoblotting. (m) Sirius Red staining of livers from *HuR*^flox/flox^ and *HuR-*HKO mice fed with an MCD for 3 weeks (n=5). *, p< 0.05. **, p< 0.01. Data represent the mean ± SEM.

### Hepatic deletion of *HuR* decreases expression of inflammation and cell death-related genes

To comprehensively compare the gene expression profiles in the livers of *HuR*-HKO and *HuR*^flox/flox^ mice fed an MCD, we performed an RNA sequencing (RNA-seq) analysis. As shown in Fig. 3a, 353 genes were upregulated and 422 genes were downregulated. Gene Ontology (GO) analysis showed that genes related to leukocyte migration, leukocyte chemotaxis, cell chemotaxis, neutrophil migration, Fc receptor signaling pathway, and positive regulation of apoptotic process were significantly decreased (Fig. 3b), whereas those associated with renal system development, kidney development, urogenital system development and steroid biosynthetic process were upregulated (Supplementary Fig. 2). qPCR analysis further confirmed these data. As shown in Fig. 3c, *Mmp9*, *Tgf1β*, *Tnfa*, *I1β*, *Il6*, *Ccl2*, *Ccl5*, *iNos*, *Cxcl2*, *Vav1*, *Csf1*, *NFκB2*, *Cd14*, *DR5* and *Casp9* mRNA levels were downregulated in *HuR*-HKO mice.

**Figure 3.**
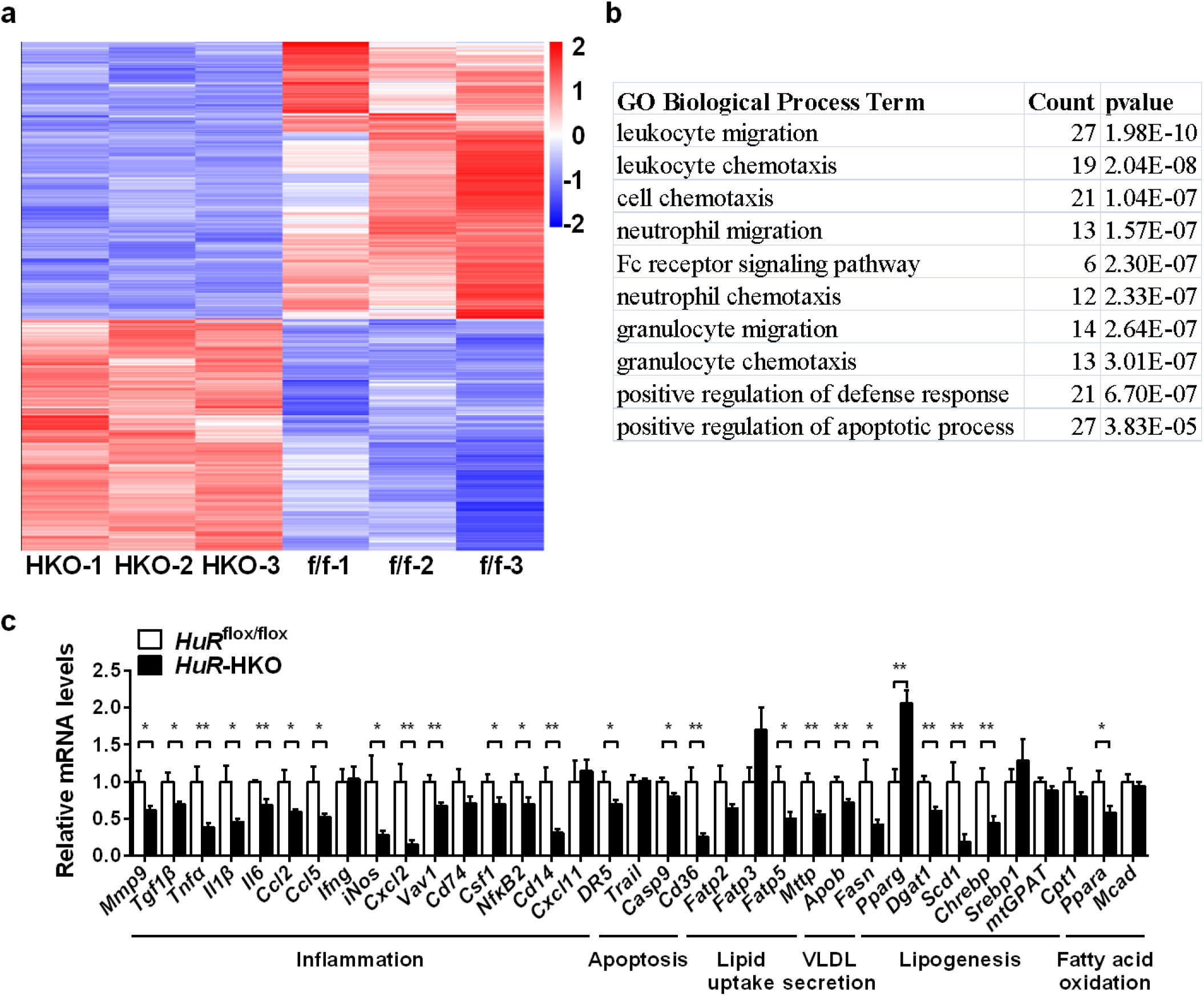
*HuR*–HKO mice display reduced expression of genes related to inflammation, cell death, and lipid metabolism compared with *HuR*^flox/flox^ mice under an MCD feeding condition. (a) RNA-seq analysis was performed in the livers of *HuR*^flox/flox^ and *HuR-*HKO mice fed with an MCD for 3 weeks (n=3). The differentially expressed genes (DEGs) (HKO VS f/f) including 422 downregulated genes and 353 upregulated genes were shown in a heatmap (|log2foldchange|>1.5 and pval<0.05). (b) Top GO biological process terms enriched in downregulated genes. (c) Real-time qPCR analysis of mRNA levels in the livers of *HuR*^flox/flox^ and *HuR-*HKO mice fed an MCD for 3 weeks (n=7-9). *, p< 0.05. **, p< 0.01. Data represent the mean ± SEM.

Hepatic lipid accumulation contributes to NASH progression ^7, 38^. Liver steaotosis is an imbalance between fatty acid uptake, lipogenesis, fatty acid β-oxidation, and VLDL secretion ^4, 39^. To determine whether fatty acid uptake, fatty acid β-oxidation, lipogenesis, or VLDL secretion contributes to the decreased liver steaotosis in *HuR*-HKO mice fed an MCD. The expression of genes related to fatty acid uptake (*Cd36*, *Fatp2*, *Fatp3* and *Fatp5*), VLDL secretion (*ApoB* and *Mttp*), lipogenesis (*Fasn*, *Pparg*, *Dgat1, Scd1*, *Chrebp*, *Srebp1*, and *mtGPAT1*), and fatty acid β-oxidation (*Cpt1α*, *Ppara,* and *Mcad*) were measured by qPCR. As shown in Fig. 3c, the expression of genes associated to VLDL secretion (*ApoB* and *Mttp*) and fatty acid β-oxidation (*Ppara*) were decreased in the livers of *HuR*-HKO mice. These data suggest that VLDL secretion and fatty acid β-oxidation less likely contributes to the decreased liver steaotosis in *HuR*-HKO mice. qPCR data also showed that genes associated to fatty acid uptake (*Cd36* and *Fatp5*) and lipogenesis (*Fasn*, *Dgat1, Scd1*, and *Chrebp*) were reduced (Fig. 3c), which may contribute to the decreased liver steaotosis in *HuR*-HKO mice under an MCD feeding condition. However, we did not observe any difference in the expression of genes related to fatty acid uptake and lipogenesis between *HuR*-HKO and *HuR*^flox/flox^ mice mice fed with a normal chow (Supplementary Fig. 3), which suggests that hepatic deletion of *HuR* does not primarily affect lipid metabolism. We consistently observed that DR5 was significantly downregulated in *HuR*-HKO mice under both normal chow and MCD feeding conditions (Fig. 2l), indicating that hepatic deletion of *HuR* primarily affects DR5 expression.

### Hepatic overexpression of *HuR* induces hepatocyte death and liver injury

Next, we asked whether hepatic overxpression of *HuR* could induce hepatocyte death and liver injury. We generated liver-specific *HuR*-overexpressing (LOE) mice were generated by injecting purified HuR adenovirus. Same amount of β Gal adenovirus injection served as Control. As expected, Flag-HuR levels were increased in the livers of *HuR-*LOE mice (Fig. 4a). Liver-specific overexpression of HuR caused liver injury, as revealed by increased blood ALT activity (Fig. 4b), increased liver weight (Fig. 4c), decreased blood glucose (Fig. 4d), increased immune cell infiltration into the liver (Fig. 4e), and increased TUNEL-positive cells (Fig. 4f). The increased cell death was likely duo to DR5/caspase 8/caspase 3 signaling pathway in the livers of *HuR-*LOE mice (Fig. 4g). The expression of DR5 was significantly increased in *HuR-*LOE mice (Fig. 4g,h). These data indicate that hepatic overexpression of *HuR* leads to hepatocyte death and liver injury, which is likely due to increased DR5 expression.

**Figure 4.**
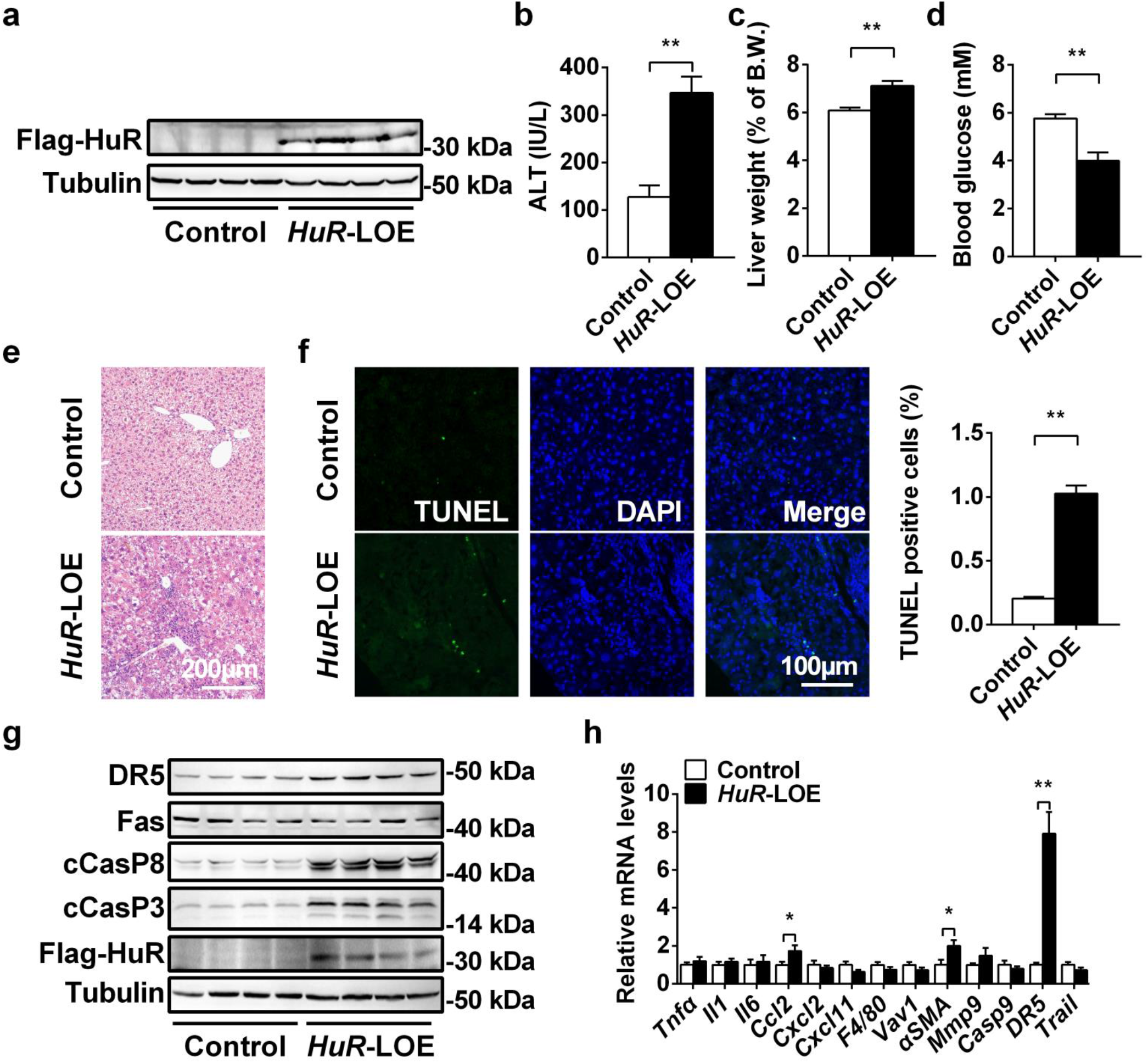
Hepatic overexpression of *HuR* induces hepatocyte death, liver inflammation and injury. (a) Flag-HuR protein levels in the livers of Control and *HuR*-LOE mice were measured by immunoblotting. (b) Serum ALT activities (n=8-9). (c) Liver weights (n=7). (d) Blood glucose levels (n=7). (e) Representative H&E staining of liver sections from Control and *HuR*-LOE mice. (f) TUNEL positive cells (n=5). (g) The death receptors (FAS and DR5) /caspase8/caspase3 signaling pathway was measured by immunoblotting in Control and *HuR*-LOE mice. (h) Relative mRNA levels (n=9) *, p< 0.05. **, p< 0.01. Data represent the mean ± SEM.

### HuR promotes cell death in primary hepatocytes by increasing both DR5 expression and the cleavage of caspase 8 and caspase 3

To determine whether HuR directly triggers hepatocyte death, we performed cell viability assay in HuR-overexpressing primary hepatocytes. The expression levels of Flag-tagged HuR were high (Fig. 5a), and the cell viability were significantly decreased (Fig. 5b). To comprehensively analyze the transcriptinal changes in the HuR-overexpressing primary hepatocytes, we performed an RNA sequencing (RNA-seq) analysis. As shown in Fig. 5c, 2677 genes were upregulated and 2712 genes were downregulated. Gene Ontology (GO) analysis showed that genes related to inflammation and apoptotic signaling pathway were significantly increased, whereas the genes related to catabolic process were dramatically decreased (Fig. 5d). qPCR analysis further confirmed these data. As shown in Fig. 5e, *TNFa*, *I1β*, *Il6*, *Ccl2*, *Cxcl2*, *Cxcl11*, *F4/80*, *Vav1*, *αSMA*, *Mmp9*, *Casp9*, and *Dr5* mRNA levels were increased in *HuR*-HKO mice.

**Figure 5.**
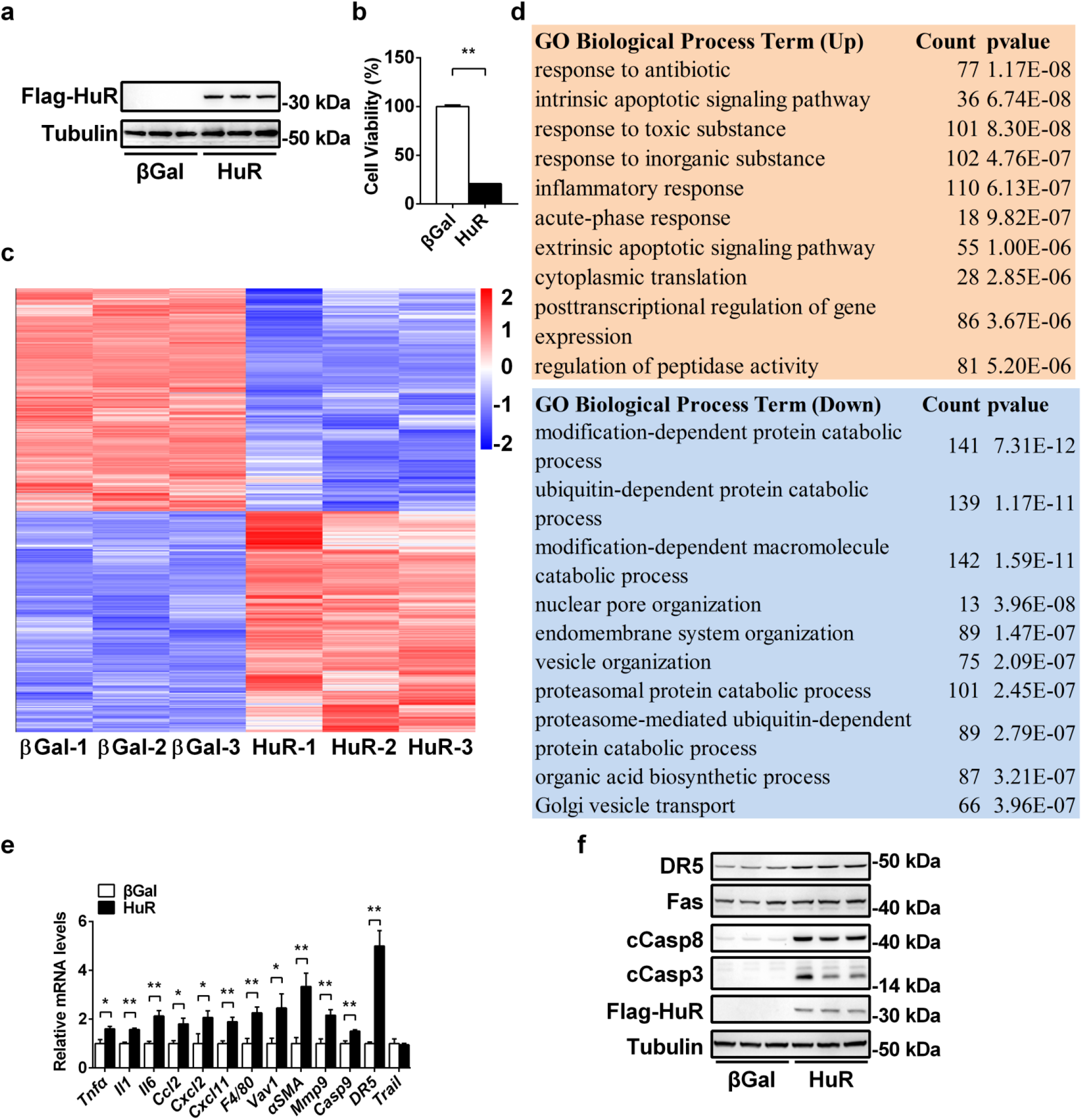
HuR promotes hepatocyte death by increasing both DR5 expression and the cleavage of caspase 8 and caspase 3. (a) Flag-HuR protein levels in the livers were measured by immunoblotting. (b) Cell viability assay in the βGal- and HuR-overexpressing hepatocytes, respectively (n=5). (c) RNA-seq analysis was performed in the βGal-and HuR-overexpressing hepatocytes (n=3). The differentially expressed genes (DEGs) (βGal VS HuR) including 2712 downregulated genes and 2677 upregulated genes were shown in a heatmap (|log2foldchange|>1.5 and pval<0.05). (d) Top GO biological process terms enriched in upregulated and downregulated genes. (e) Real-time qPCR analysis of mRNA levels in the βGal-and HuR-overexpressing hepatocytes. (n=3-6). (f) The death receptors (FAS and DR5) /caspase8/caspase3 signaling pathway was measured by immunoblotting. *, p< 0.05. **, p< 0.01. Data represent the mean ± SEM.

To determine whether this cell death induced by HuR is associated with death receptors/caspase8/caspase 3 pathway, we measured death receptor protein levels and the cleavage of caspase 8 and caspase 3. As shown in Fig. 5f, the DR5 protein levels and the cleavage of caspase 8 and caspase 3 were significantly increased in HuR-overexpressing hepatocytes. We did not observe any difference in FAS protein levels between βGal and HuR groups (Fig. 5f), indicating that HuR does not regulate FAS expression. TRAIL, the ligand for DR5, was not regulated by HuR (Fig. 5e). These data demonstrate that HuR induces hepatocyte death by enhancing DR5/caspase8/caspase3 pathway.

### HuR is essential for PA&TNFα-induced cell death in primary hepatocytes

A combination of PA and TNFα (PA&TNFα) can induce hepatocyte death that mimics NASH-associated hepatocyte death. To determine whether HuR is essential for PA&TNFα-induced hepatocyte death, we isolated primary hepatocytes from *HuR*-HKO and *HuR*^flox/flox^ mice and treated these cells with PA&TNFα. As shown in Fig. 6a, HuR deficiency protected hepatocytes from PA&TNFα-induced cell death. Expectedly, HuR and DR5 protein levels were increased by PA&TNFα treatment in *HuR*^flox/flox^ hepatocytes, whereas their protein levels were significantly decreased in *HuR*-HKO hepatocytes (Fig. 6b). Moreover, the cleavage of caspase 8 and caspase 3 was increased by PA&TNFα treatment in *HuR*^flox/flox^ hepatocytes, but their cleavage was significantly blocked in *HuR*-HKO hepatocytes (Fig. 6b). The ligand for DR5 is TRAIL. To determine whether HuR is essential for TRAIL-induced activation of caspase8 and caspase3, primary hepatocytes were isolated from *HuR*-HKO and *HuR*^flox/flox^ mice, and TRAIL-induced cleavage of caspase8 and caspase3 was measured by immunoblotting. As shown in Fig. 6c, the cleavage of caspase 8 and caspase 3 were significantly decreased in *HuR*-HKO hepatocytes. Consistently, DR5 protein levels were significantly decreased in *HuR*-HKO hepatocytes (Fig. 6b,c). These data demonstrate that HuR is essential for PA&TNFα-induced hepatocyte death and the cleavage of caspase8 and caspase 3, most likely due to the decreased DR5 protein levels.

**Figure 6.**
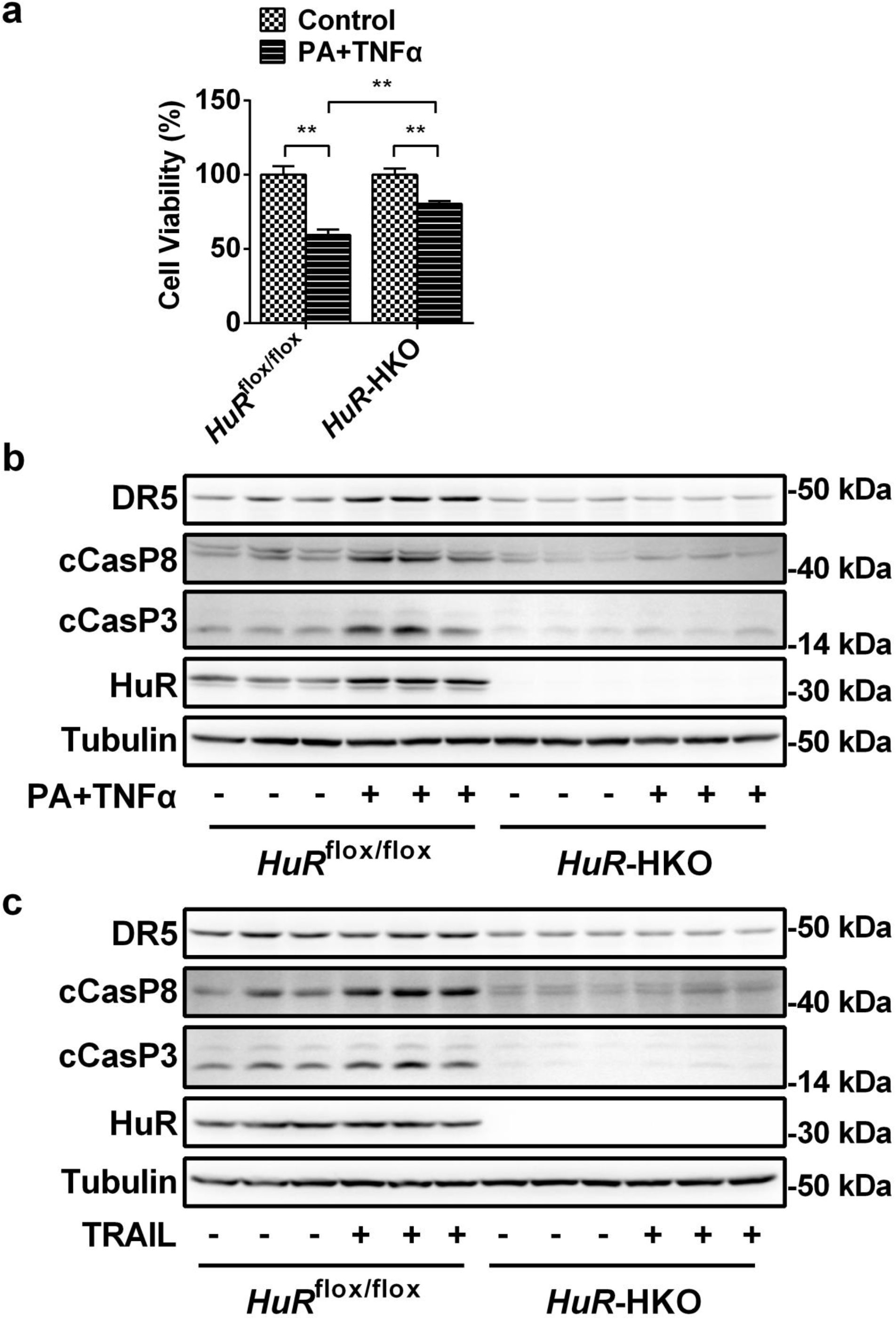
HuR is essential for maintaining DR5/caspase8/caspase3 signaling pathway in primary hepatocytes. (a) Primary hepatocytes were isolated from *HuR*-HKO and *HuR*^flox/flox^ mice and treated with PA (1 mM) and TNFα (100 ng/ml) for 28 hours. Cell viability assay was measured by MTT. (b) Primary hepatocytes were isolated from *HuR*-HKO and *HuR*^flox/flox^ mice and treated with PA (1 mM) and TNFα (100g/ml) for 24 hours. The cleavage of caspase8 and caspase3 was measured by immunoblotting. (c) Primary hepatocytes were isolated from *HuR*-HKO and *HuR*^flox/flox^ mice and treated with TRAIL (1μg/ml) for 2 hours. The cleavage of caspase8 and caspase3 was measured by immunoblotting. These experiments were at least repeated twice with similar results. *, p< 0.05. **, p< 0.01. Data represent the mean ± SEM.

### HuR directly binds to the 3’-UTR of DR5 transcript and decreases its mRNA decay

HuR has been shown to binds to AREs in 3’-UTR and regulate mRNA stability. Photoactivatable ribonucleoside-enhanced cross-linking and immunoprecipitation (PAR-CLIP) sequencing showed that HuR binds to DR5 transcript ^40^. RNA immunoprecipitation (RIP) sequencing analysis also showed that HuR binds to DR5 (TNFRSF10B) mRNA ^41^. Sequence analysis showed that 3’-UTR of DR5 mRNA contains several AREs, which further indicates that HuR may bind to 3’-UTR of DR5 mRNA and regulate DR5 mRNA stability. To test this hypothesis, we performed RIP-RT-qPCR analysis in both HuR-overexpressing and HuR knockout hepatocytes. The successful HuR precipitation by both Flag-beads and HuR antibody was confirmed by immunoblotting (Fig. 7a,b). As expected, both HuR and Flag immunoblotting signals were observed in *HuR*^flox/flox^ hepatocytes and Flag-HuR-overexpressing hepatocytes, respectively, whereas no immunoblotting signal of HuR in immunoprecipitation or immunoblotting was observed in HuR deficient hepatocytes (Fig. 7a,b). RIP-RT-qPCR showed that a huge increase of DR5 mRNAs retrieved by Flag-beads in Flag-HuR-overexpressing hepatocytes and also observed a significant reduction of DR5 mRNA retrieved by HuR antibody in HuR-HKO hepatocytes (Fig. 7c,d). To examine the effect of HuR on DR5 mRNA stability in hepatocytes, we used Actinomycin D to inhibit mRNA transcription in HuR-HKO and HuR-overexpressing hepatocytes and measured the decay rate for DR5 mRNA. The half-life of DR5 mRNA was increased from 5.77 to 7.66 h in the HuR-overexpressing hepatocytes (Fig. 7e), while the half-life of DR5 mRNA was decreased from 7.22 to 5.14 h in the HuR-HKO hepatocytes (Fig. 7f). Taken together, HuR may induce hepatocyte death by binding and stabilizing DR5 mRNAs, leading to the activation of caspase8/caspase3 signaling pathway.

**Figure 7.**
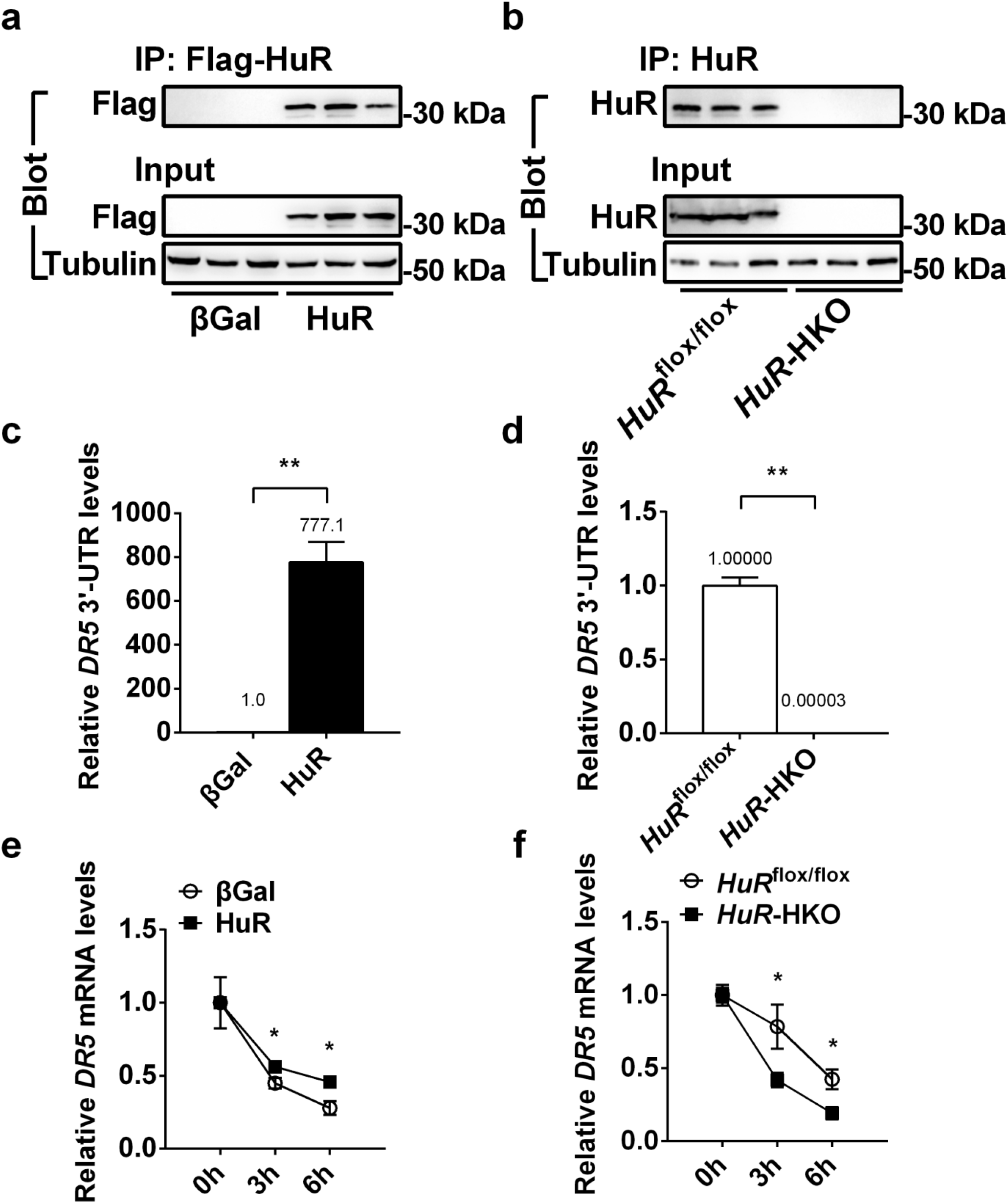
HuR directly binds to the 3’-UTR of *DR5* transcript and increases its mRNA stability. (a,b) The successful HuR immunoprecipitation by both Flag-beads and HuR antibody, respectively, was confirmed by immunoblotting. (c,d) RIP-RT-qPCR showed that a huge increase of *DR5* 3’-UTR retrieved by Flag-beads in Flag-HuR-overexpressing hepatocytes and that a significant reduction of *DR5* 3’-UTR retrieved by HuR antibody also observed in *HuR*-HKO hepatocytes (n=3). (e) RNA stability assay in the βGal- and HuR-overexpressing hepatocytes (n=6). (f) RNA stability assay in the *HuR*^flox/flox^ and *HuR-*HKO hepatocytes (n=6). *, p< 0.05. **, p< 0.01. Data represent the mean ± SEM.

## Discussion

Hepatocyte death is at the center of the second hits during NASH pathogenesis because it triggers liver inflammation, liver injury, and fibrosis. Clinical studies and genetic manipulations demonstrate that hepatocyte apoptosis plays a key role in NASH progression ^8, 9, 10, 11, 12^. However, whether RNA processing regulates death signaling pathway during NASH progression remain poorly understood. In this study, we demonstrated that abnormal elevated HuR promotes hepatocyte apoptosis, inflammation and diet-induced NASH in mice by stabilizing DR5 mRNA. Hepatic deletion of HuR ameliorates MCD-induced NASH by decreasing DR5/caspase 8/caspase 3 signaling pathway, which provides a new therapeutic target for the treatment of NASH in humans.

Several lines of evidence support the pathological significance of HuR in controlling NASH progression. Hepatic HuR, especially the cytosolic HuR, is abnormally upregulated in both human patients with NASH and mice with MCD-induced NASH. PA and TNFα could increased both HuR expression and cytosolic localization of HuR in primary hepatocytes, which may be partially due to PKC-mediated phosphorylation of HuR ^33^. Hepatic deletion of *HuR* ameliorates the severity of diet-induced NASH in mice, as shown by the reduced serum ALT activity, the decreased hepatic steatosis, the decreased hepatocyte death, and the reduced liver inflammation. In contrast, liver-specific overexpression of HuR induces hepatocyte death and liver injury. HuR overexpression induced hepatocyte death *in vitro* in a cell-autonomous manner, demonstrating a direct role of HuR in triggering liver damage.

HuR regulates biological and pathophysiological processes by targeting and stabilizing ARE-enriched transcripts ^42, 43^. HuR overexpression in primary hepatocytes causes hepatocyte death by increasing ARE-enriched transcripts, which includes apoptosis and inflammation-related transcripts. One of the most robust response transcripts is *DR5*. DR5 activation directly triggers caspase 8 cleavage to induce its activation and further increases the cleavage of caspase 3, leading to apoptosis ^37^. DR5 plays a key role in fatty acid-induced cell death and is also induced during NASH pathogenesis ^9, 10^. Knockdown of *DR5* ameliorates fatty acid-induced hepatocyte death ^9^. However, the pathophysiological cues that impinge on DR5 remain largely unknown. Our current work uncovers *DR5* mRNA stability mediated by HuR as a key regulator of NASH progression. HuR directly binds to the 3’-UTR of *DR5* mRNA, and then strongly increases DR5 expression, which leads to the activation of caspase 8 and caspase 3 in hepatocytes, resulting in the increased hepatocyte death and liver injury. More importantly, HuR is essential for DR5 expression in the liver. Hepatic deletion of *HuR* decreases the expression of DR5 in the isolated primary hepatocytes and the livers of mice fed with either a normal chow diet or an MCD diet, which is most likely due to the decreased RNA stability of DR5 transcript. HuR deficiency ameliorates PA&TNFα-induced hepatocyte death, which is likely due to decreased both DR5 expression and the activation of caspase8/caspase3 signaling pathway. These results demonstrate that HuR is essential for the activation of DR5/caspase8/caspase3 signaling pathway in hepatocytes.

Although some lipid metabolism-related genes were downregulated in *HuR*-HKO mice fed with an MCD, no big difference was observed in lipid metabolism between *HuR*-HKO and *HuR*^flox/flox^ mice fed with a normal chow. Overexpression of HuR does not increase the expression of genes related to lipid metabolism. These results suggest that HuR does not primarily regulate lipid metabolism.

In conclusion, we have demonstrated that HuR is a novel important regulator of NASH progression. Our data also reveal a new mechanism in which HuR increases the mRNA stability of *DR5*. Abnormally elevated HuR expression leads to increased mRNA stability of *DR5*, which further increases hepatocyte death, contributing to NASH progression. Our results also indicate that HuR may serve as a novel drug target for the treatment of NASH.

## Acknowledgements

This study was supported by the National Natural Science Foundation of China Grant (31671225 and 31971083) and Natural Science Foundation of Heilongjiang Province (YQ2019C011). X.L. is supported by Chinese Postdoctoral Science Foundation (AUGA4130900619).

## Author contributions

X.L. performed most of the experiments. B.Y. initiated this project and collected part of the key data. Z.Y. provided human liver samples and researched data. X.S. performed part of qPCR experiments. W.G. provided *HuR*^flox/flox^ mice and reviewed manuscript. Z.C. conceived and designed the project, researched data, and wrote manuscript.

## References

1. Diehl AM, Day C. Cause, Pathogenesis, and Treatment of Nonalcoholic Steatohepatitis. New England Journal of Medicine 2017, 377(21):2063–2072.

2. Anstee QM, Reeves HL, Kotsiliti E, Govaere O, Heikenwalder M. From NASH to HCC: current concepts and future challenges. Nat Rev Gastroenterol Hepatol 2019, 16(7):411–428.

3. Day CP, James OF. Steatohepatitis: a tale of two “hits”? Gastroenterology 1998, 114(4):842–845.

4. Friedman SL, Neuschwander-Tetri BA, Rinella M, Sanyal AJ. Mechanisms of NAFLD development and therapeutic strategies. Nat Med 2018, 24(7):908–922.

5. Donnelly KL, Smith CI, Schwarzenberg SJ, Jessurun J, Boldt MD, Parks EJ. Sources of fatty acids stored in liver and secreted via lipoproteins in patients with nonalcoholic fatty liver disease. J Clin Invest 2005, 115(5):1343–1351.

6. Schuster S, Cabrera D, Arrese M, Feldstein AE. Triggering and resolution of inflammation in NASH. Nat Rev Gastroenterol Hepatol 2018, 15(6):349–364.

7. Sanyal AJ. Past, present and future perspectives in nonalcoholic fatty liver disease. Nat Rev Gastroenterol Hepatol 2019, 16(6):377–386.

8. Akazawa Y, Nakao K. To die or not to die: death signaling in nonalcoholic fatty liver disease. J Gastroenterol 2018, 53(8):893–906.

9. Cazanave SC, Mott JL, Bronk SF, Werneburg NW, Fingas CD, Meng XW, et al. Death Receptor 5 Signaling Promotes Hepatocyte Lipoapoptosis. Journal of Biological Chemistry 2011, 286(45):39336–39348.

10. Farrell GC, Larter CZ, Hou JY, Zhang RH, Yeh MM, Williams J, et al. Apoptosis in experimental NASH is associated with p53 activation and TRAIL receptor expression. Journal of gastroenterology and hepatology 2009, 24(3):443–452.

11. Guo L, Zhang P, Chen Z, Xia H, Li S, Zhang Y, et al. Hepatic neuregulin 4 signaling defines an endocrine checkpoint for steatosis-to-NASH progression. The Journal of Clinical Investigation 2017, 127(12):4449–4461.

12. Hatting M, Zhao G, Schumacher F, Sellge G, Al Masaoudi M, Gaβler N, et al. Hepatocyte caspase-8 is an essential modulator of steatohepatitis in rodents. Hepatology (Baltimore, Md) 2013, 57(6):2189–2201.

13. Feldstein AE, Canbay A, Angulo P, Taniai M, Burgart LJ, Lindor KD, et al. Hepatocyte apoptosis and fas expression are prominent features of human nonalcoholic steatohepatitis. Gastroenterology 2003, 125(2):437–443.

14. Hirsova P, Ibrahim SH, Krishnan A, Verma VK, Bronk SF, Werneburg NW, et al. Lipid-Induced Signaling Causes Release of Inflammatory Extracellular Vesicles From Hepatocytes. Gastroenterology 2016, 150(4):956–967.

15. Machado MV, Michelotti GA, Pereira TdA, Boursier J, Kruger L, Swiderska-Syn M, et al. Reduced lipoapoptosis, hedgehog pathway activation and fibrosis in caspase-2 deficient mice with non-alcoholic steatohepatitis. Gut 2015, 64(7):1148–1157.

16. Hirsova P, Weng P, Salim W, Bronk SF, Griffith TS, Ibrahim SH, et al. TRAIL Deletion Prevents Liver, but Not Adipose Tissue, Inflammation during Murine Diet-Induced Obesity. Hepatology communications 2017, 1(7):648–662.

17. Witek RP, Stone WC, Karaca FG, Syn WK, Pereira TA, Agboola KM, et al. Pan-caspase inhibitor VX-166 reduces fibrosis in an animal model of nonalcoholic steatohepatitis. Hepatology (Baltimore, Md) 2009, 50(5): 1421–1430.

18. Barreyro FJ, Holod S, Finocchietto PV, Camino AM, Aquino JB, Avagnina A, et al. The pan-caspase inhibitor Emricasan (IDN-6556) decreases liver injury and fibrosis in a murine model of non-alcoholic steatohepatitis. Liver international: official journal of the International Association for the Study of the Liver 2015, 35(3):953–966.

19. Suresh Babu S, Joladarashi D, Jeyabal P, Thandavarayan RA, Krishnamurthy P. RNA-stabilizing proteins as molecular targets in cardiovascular pathologies. Trends in Cardiovascular Medicine 2015, 25(8):676–683.

20. Pascale A, Govoni S. The complex world of post-transcriptional mechanisms: is their deregulation a common link for diseases? Focus on ELAV-like RNA-binding proteins. Cell Mol Life Sci 2012, 69(4):501–517.

21. Hinman MN, Lou H. Diverse molecular functions of Hu proteins. Cellular and Molecular Life Sciences 2008, 65(20):3168.

22. Qin W, Li X, Xie L, Li S, Liu J, Jia L, et al. A long non-coding RNA, APOA4-AS, regulates APOA4 expression depending on HuR in mice. Nucleic Acids Research 2016, 44(13):6423–6433.

23. Vazquez-Chantada M, Fernandez-Ramos D, Embade N, Martinez-Lopez N, Varela-Rey M, Woodhoo A, et al. HuR/methyl-HuR and AUF1 regulate the MAT expressed during liver proliferation, differentiation, and carcinogenesis. Gastroenterology 2010, 138(5):1943–1953.

24. Woodhoo A, Iruarrizaga-Lejarreta M, Beraza N, Garcia-Rodriguez JL, Embade N, Fernandez-Ramos D, et al. Human antigen R contributes to hepatic stellate cell activation and liver fibrosis. Hepatology (Baltimore, Md) 2012, 56(5):1870–1882.

25. Ale-Agha N, Galban S, Sobieroy C, Abdelmohsen K, Gorospe M, Sies H, et al. HuR regulates gap junctional intercellular communication by controlling beta-catenin levels and adherens junction integrity. Hepatology (Baltimore, Md) 2009, 50(5):1567–1576.

26. Ghosh M, Aguila HL, Michaud J, Ai Y, Wu MT, Hemmes A, et al. Essential role of the RNA-binding protein HuR in progenitor cell survival in mice. J Clin Invest 2009, 119(12):3530–3543.

27. Wang Y, Guo Y, Tang C, Han X, Xu M, Sun J, et al. Developmental Cytoplasmic-to-Nuclear Translocation of RNA-Binding Protein HuR Is Required for Adult Neurogenesis. Cell reports 2019, 29(10):3101–3117.e3107.

28. Ren X, Li X, Jia L, Chen D, Hou H, Rui L, et al. A small-molecule inhibitor of NF-κB-inducing kinase (NIK) protects liver from toxin-induced inflammation, oxidative stress, and injury. The FASEB Journal 2017, 31(2):711–718.

29. Jia L, Jiang Y, Li X, Chen Z. Purbeta promotes hepatic glucose production by increasing Adcy6 transcription. Mol Metab 2020, 31:85–97.

30. Li X, Jia L, Chen X, Dong Y, Ren X, Dong Y, et al. Islet α-cell Inflammation Induced By NF-κB inducing kinase (NIK) Leads to Hypoglycemia, Pancreatitis, Growth Retardation, and Postnatal Death in Mice. Theranostics 2018, 8(21):5960–5971.

31. Wang Y, Gao M, Zhu F, Li X, Yang Y, Yan Q, et al. METTL3 is essential for postnatal development of brown adipose tissue and energy expenditure in mice. Nature Communications 2020, 11(1):1648.

32. Liu J, Eckert MA, Harada BT, Liu SM, Lu Z, Yu K, et al. m(6)A mRNA methylation regulates AKT activity to promote the proliferation and tumorigenicity of endometrial cancer. Nature cell biology 2018, 20(9):1074–1083.

33. Lu S, Mott JL, Harrison-Findik DD. Saturated fatty acids induce post-transcriptional regulation of HAMP mRNA via AU-rich element-binding protein, human antigen R (HuR). The Journal of biological chemistry 2015, 290(40):24178–24189.

34. Martinez-Chantar ML, Vazquez-Chantada M, Garnacho M, Latasa MU, Varela-Rey M, Dotor J, et al. S-adenosylmethionine regulates cytoplasmic HuR via AMP-activated kinase. Gastroenterology 2006, 131(1):223–232.

35. Ricchi M, Odoardi MR, Carulli L, Anzivino C, Ballestri S, Pinetti A, et al. Differential effect of oleic and palmitic acid on lipid accumulation and apoptosis in cultured hepatocytes. Journal of gastroenterology and hepatology 2009, 24(5):830–840.

36. Kudo H, Takahara T, Yata Y, Kawai K, Zhang W, Sugiyama T. Lipopolysaccharide triggered TNF-α-induced hepatocyte apoptosis in a murine non-alcoholic steatohepatitis model. Journal of Hepatology 2009, 51(1):168–175.

37. Malhi H, Guicciardi ME, Gores GJ. Hepatocyte Death: A Clear and Present Danger. Physiological reviews 2010, 90(3):1165–1194.

38. Machado MV, Diehl AM. Pathogenesis of Nonalcoholic Steatohepatitis. Gastroenterology 2016, 150(8):1769–1777.

39. Kawano Y, Cohen DE. Mechanisms of hepatic triglyceride accumulation in non-alcoholic fatty liver disease. J Gastroenterol 2013, 48(4):434–441.

40. Kishore S, Jaskiewicz L, Burger L, Hausser J, Khorshid M, Zavolan M. A quantitative analysis of CLIP methods for identifying binding sites of RNA-binding proteins. Nature Methods 2011, 8(7):559–564.

41. Bersani C, Huss M, Giacomello S, Xu L-D, Bianchi J, Eriksson S, et al. Genome-wide identification of Wig-1 mRNA targets by RIP-Seq analysis. Oncotarget 2016, 7(2):1895–1911.

42. Mukherjee N, Corcoran DL, Nusbaum JD, Reid DW, Georgiev S, Hafner M, et al. Integrative regulatory mapping indicates that the RNA-binding protein HuR couples pre-mRNA processing and mRNA stability. Molecular cell 2011, 43(3):327–339.

43. Lebedeva S, Jens M, Theil K, Schwanhäusser B, Selbach M, Landthaler M, et al. Transcriptome-wide analysis of regulatory interactions of the RNA-binding protein HuR. Molecular cell 2011, 43(3):340–352.

